# Over Two Million Protein Translocations Reveal Optimal Conditions for High Bandwidth Recordings, Studying Transport Energetics, and Unfolding

**DOI:** 10.1101/2022.09.28.509929

**Authors:** Y. M. Nuwan, D. Y. Bandara, Kevin J. Freedman

## Abstract

The gradual tapered geometry of nanopipettes offers a unique perspective on protein transport through nanopores since both a gradual and fast confinement is possible depending on translocation direction. Protein capture rate, unfolding, speed of translocation, and clogging probability are studied by toggling the lithium chloride concentration between 4 M and 2 M. Interestingly, the proteins in this study could be transported with or against the electrophoresis and offer vastly different attributes of sensing and affect whether a protein unfolds during pore transit. A ruleset for studying proteins is developed that prevents irreversible pore clogging and yielded upwards of >100,000 events/nanopore. Minimizing clogging also permitted higher quality data via the use of smaller pores (i.e., <2× the size of the protein) including higher SNR recordings and data acquisition at the highest available bandwidth (100 kHz). The extended duration of experiments further revealed that the capture rate takes ~2 hours to reach a steady state with a value ~3x greater than the initial reading, emphasizing the importance of reaching equilibrated transport for studying the energetics of protein transport (i.e., diffusion vs barrier-limited). Even in the equilibrated transport state, improper lowpass filtering was shown to distort the classification of diffusion-limited vs barrier-limited transport. Finally electric-field induced protein unfolding was found to be most prominent in EO dominant transport whereas EP dominant events show no evidence of unfolding. Thus, our findings showcase the optimal conditions for protein translocations and the impact on studying protein unfolding, transport energetics, and acquiring high bandwidth data.

## Introduction

Nanopores are often perceived as a sensor class capable of delivering single molecule-level information at an unprecedented throughput at a fraction of the cost compared to most contemporary analytical methods and boasts prospects as far as protein and DNA sequencing. ^1–2^ The application scope of nanopores is vast and spans a range of biomolecules and bioparticles.^3–7^ Fast protein translocations often challenge the bandwidth of conventional electronics (typically ≤100 kHz) leading to attenuated electrical readouts.^8^ Advancements in electronics have led to more compact high bandwidth instruments that are capable of circumventing signal attenuation^9^ while high open-pore noise precludes its application with most membrane materials. Optimizing the sensing conditions to improve the signal-to-noise ratio (SNR) would pave the way for the widespread adoption of high bandwidth acquisition approaches. A rich blend of approaches to slow down the translocation speed of analytes have been proposed over the years, including, but not limited to pressure opposition,^10^ surface coating and anchoring and,^6, 11^ surfaces capturing. These are well complemented by efforts for noise reduction through membrane coatings (e.g., PDMS),^12^ surface passivation,^13^ low-noise material (e.g., quartz),^14^ and amplifiers (e.g., CMOS preamplifiers)^9^. As such, understanding the intricate links of protein translocation characteristics (e.g., SNR, speed) with chemical (e.g., solution and pore-surface and membrane chemistry), physical (e.g., pore size, membrane thickness), and electronic (e.g., bandwidth) conditions are imperative to achieve desirable sensing conditions for proteins.

Compared to the plethora of work done with DNA, the protein footprint in nanopipette studies is meager. A major hurdle with proteins is the nonspecific adsorption leading to irreversible pore clogging which is often more profound than that observed with DNA. Refuting pore clogging is challenging and often accepted as part of nanopore experiments which precludes the operation of pores over longer periods and the collection of large amounts of data (i.e., tens of thousands instead of a few hundred to few thousand). This is very critical for nanopores to venture into the commercial space since the pore should be open for *business* rather than succumbing to analyte clogging in a short time frame. Refuting analyte clogging would be one aspect that would be discussed in this work. Another important aspect of protein (or any other analyte) translocations is the SNR. Proteins are often translocated in the electrophoretic direction, and it is not uncommon to see the use of high salt conditions mainly due to the poor SNR associated with protein translocations under low salt conditions. For a negatively charged protein, the electroosmotic transport originating from negatively charged pore material (e.g., quartz, silicon nitride) has an added advantage of opposing electrophoretic and electroosmotic forces that can slow down the translocation speed of the protein. One way to increase SNR would be to use a smaller pore. However, this increases the irreversible sticking probability rendering the pore futile for further use. As shown later, the tapered geometry of nanopipettes provides a solution to this in which we observed pipette to bath transport (i.e., forward translocations) is less prone to clogging compared to bath to pipette transport (i.e., backward translocations). The geometry may be preventing/discouraging the co-translocation of proteins in the forward direction whereas such constraints are not present in the backward direction (analogous to planar nanopore configuration). Thus, the forward direction permits the use of smaller pores to improve SNR, and by extension, as shown in the manuscript, allows the maximum available lowpass filter (LPF) of the Axopatch 200B (i.e., 100 kHz) to be used to detect protein translocations (with four different proteins) across a broad range of voltages (i.e., ±300 mV to ±1000 mV).

We used the holo-form of the human serum transferrin (hSTf) as the core model protein in this study. It’s a blood glycoprotein protein with a molar mass of ~80 kDa and a pI of ~5.1-5.5 which is critical for iron transport.^15^ The transport properties of hSTf with planar nanopores are well-characterized,^16–18^ thus providing a frame of reference when required. In this study, we investigated a wide range of tunable parameters to showcase the dependence of the transport properties (e.g., SNR, translocation time, and mode of transport (i.e., electroosmosis (EO) or electrophoresis (EP) dominant)) and their implications on protein structure (i.e., voltage-driven unfolding). More broadly, the nanopipette size, solution chemistry, and transport direction were investigated broadly with a range of proteins. More specifically, 20 different pore sizes spanning ~9 nm to ~30 nm in the electroosmotic realm (2M LiCl), 5 different pore sizes spanning ~7 nm to ~12.5 nm in the electrophoretic realm (4M LiCl), and, 8 different voltages in most instances (±300 mV to ±1000 mV in ±100 mV increments) were investigated for a total event count of 2.8 million protein translocations (see Supporting Information). The transport direction dependence was investigated using four proteins: hSTf, bovine serum albumin (BSA), ferritin, and hemoglobin. The isoelectric points for BSA, hemoglobin (bovine) and, ferritin are reported to be ~4.5, ~7.1 and, ~5.4-5.5 respectively. As mentioned previously, the forward direction is more conducive for events (often producing >20,000 events/pipette and even >100,000 events/pipette) while the backward direction more often leads to irreversible clogging. Furthermore, smaller pore sizes (<2× the size of proteins) can be used with relative ease in the forward direction which improves volumetric occlusion by the protein, confinement, and by extension, SNR enabling high LPF experiments that are crucial for temporally non-attenuated protein sensing. Thus, these findings are broadly beneficial to exploring protein sensing applications and biophysics.

## Materials and Methods

### Nanopipette Fabrication

Quartz nanopipettes (QF10070-7.5, Sutter Instruments) were fabricated using Sutter P-2000 with the following settings: Heat of 630, Fil of 4, Velocity of 61, Delay of 130. The pull was varied from 125-220 to tune the final diameter of the nanopore (see Figure 1 and SI Figure S1 for the relationship of pull vs pore diameter). The following equation was used to estimate the nanopore diameter (*d_p_*) which was derived previously^19^:

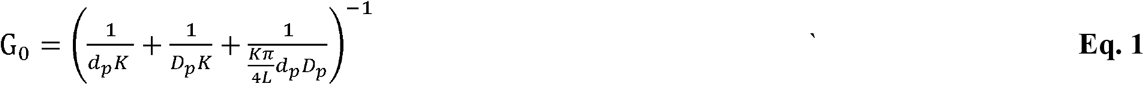

where G_0_, K, *D_p_*, L, are open-pore conductance, the conductivity of the electrolyte, base diameter, and sensing length respectively.

**Figure 1:**
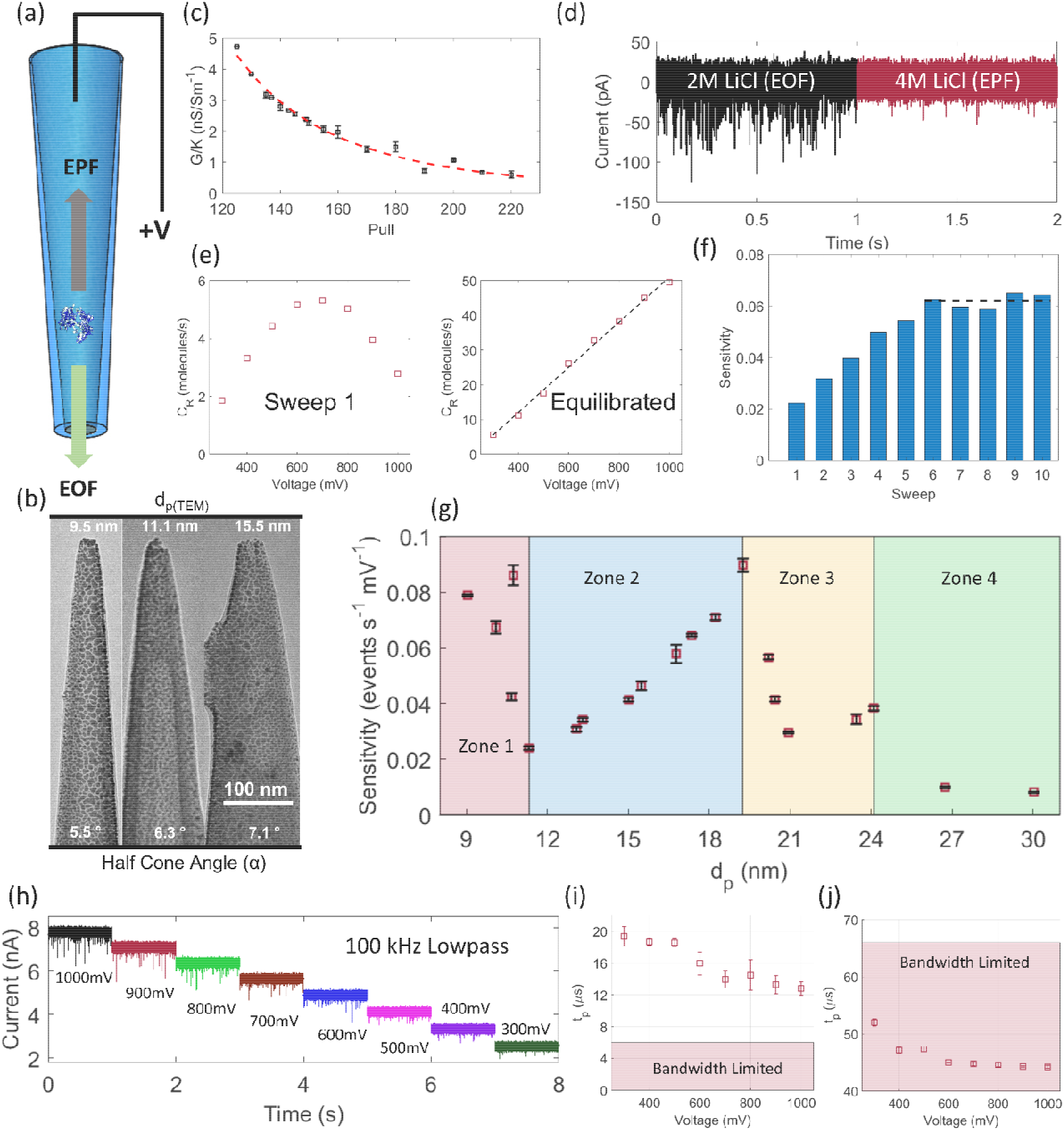
**(a)** Nanopipette setup wherein the protein is added to the pipette and driven either electroomostically or electrophoretically through the tapered nanopore. Green and gray arrows show the electroosmotic force (EOF) and electrophoretic force (EPF) directions. **(b)** TEM images of nanopipettes fabricated with a pull value of 210, 180, and 150 with an estimated diameter of ( ) ~9.5 nm, ~11.1 nm, and ~15.5 nm respectively. **(c)** The relationship between pull and conductance normalized with solution conductivity (G/K). The raw data were fitted with a function in the form *P* = (*G/K*)^*n*^ (red trace, see main text). **(d)** A concatenated trace consisting of 1-second representative traces (through a ~20 nm diameter pore) in 2M LiCl (black) and 4M LiCl (maroon). **(e)** The repercussion of poor equilibration in which a diffusionlimited transport resembles a scenario where events are missed due to fast translocations. Therefore, multiple capture rate (*C_R_*) vs voltage curves are obtained (i.e., sweeps) as shown in figure **(f),**and the mean of the plateaued region (black dashed line) is considered as the final sensitivity of the experiment. **(g)** Sensitivity vs pore size (*d_p_*, calculated from equation 1) for pores fabricated from pulls ranging from 125 to 220. The sensitivity of the largest pore was divided by two to account for the twice higher concentration used to obtain the data. **(h)** Representative current trace of a ~9 nm pore in zone-1 lowpass filtered at 100 kHz. Corresponding peak translocation time (*τ_p_*) at **(i)** 100 kHz and **(j)** 10 kHz LPF settings. All translocation-based experiments were performed at pH ~8

### Biomolecule Preparation

Stock solutions of hSTf were prepared by dissolving the as-supplied product (616397, Sigma Aldrich) in 1 mM KHCO_3_ (P235, Fisher Scientific). It was then added to the electrolyte to a final concentration of ~600nM. Hemoglobin (J63838.06, Alfa Aesar) was prepared by dissolving the as-supplied product in water. BSA (B9000S, New England Biolabs), and ferritin (S6272, MP Biomedicals) were used as supplied with appropriate dilutions. All stock solutions except BSA were stored at 4°C (BSA was stored at −20 °C).

### Electrolyte Preparation

LiCl (310468, Sigma Aldrich) was dissolved in ultra-pure water and buffered with 10 mM tris-EDTA buffer (BP2475, Fisher Scientific). The pH was adjusted by adding concentrated drops of HCl (SA48-1, Fischer Scientific) or KOH (LC192402, LabChem) and measured using Accumet AB200 pH meter.

### Electrical Measurements

Axopatch 200B (Molecular Devices LLC) connected to a Digidata 1550B (Molecular Devices LLC) was used for all measurements. Signals were filtered using the inbuilt Bessel lowpass filter of the Axopatch 200B (10 kHz or 100 kHz) and acquired at 250 kHz (500 kHz in the case of 100 kHz lowpass filter) using the digitizer. Data were then extracted using the *EventPro* app.^20^

## Results and Discussion

### Equilibration of Protein Transport and Bandwidth Limitations

Nanopipettes were fabricated conveniently through laser pulling and the relationship between pull and *G/K* is shown in Figure 1c for quartz capillaries with an internal diameter (ID) of 0.7 mm. The raw data were fitted with a function (only as a guide to the eye) in the form *P* = (*G/K*)^*n*^ where *P* and *n* are pull and an arbitrary exponent for fitting purposes (~3.6) respectively. We presented the data in *G/K* instead of *d_p_* to eliminate any model-specific limitations used to calculate the pore diameter. Figure S1 shows a pull vs *G/K* plot for quartz capillaries with an ID of 0.5 mm (*n* ~4.4). Looking at the pull vs *G/K* plots corresponding to 0.7 mm ID (Figure 1c) and 0.5 mm ID (Figure S1), a given pull produces a smaller pore with the smaller ID capillary. This is in good agreement with the observations of Sun *et al* where pores as small as ~5nm were fabricated with 0.3mm ID capillaries.^21^ All experiments were carried out using LiCl electrolyte and the translocations were in the EO direction in 2M LiCl whereas it was in the EP direction in 4M LiCl. Since both the quartz pore surface and the hSTf protein (hSTf) are net-negatively charged at the operation pH (~8), the EO and EP forces would be opposing at any given voltage bias. When hSTf is added to the pipette and a voltage bias is applied to the same side, the sign of the voltage bias that would lead to translocations would shed insight into the dominant translocation mechanism. For example, when a negative potential is applied, the EP force imparted on the protein would be from the pipette to the bath side while the EO would be from the bath to the pipette side. Thus, for translocations to be feasible, EP should dominate EO. On the other hand, if a positive voltage bias is applied, the directions of EP and EO would reserve and for translocations to happen, EO should dominate EP. Since hSTf translocate under a positive bias in 2M LiCl, EO would be dominant over EP, and in 4M LiCl, since it translocate under a negative bias, EP should be dominant over EO. For the model protein hSTf, electroosmotic transport was preferred due to the comparatively poor SNR of electrophoretic transport especially in larger pores as shown in Figure 1d (*d_p_*~20 nm). Thus, with electroosmotic transport, proteins could be probed over a wide range of pore sizes compared to electrophoretic transport.

The capture rate vs voltage curves (*C_R_* – V) curves shed insight into the transport mechanism with a linear relationship indicating diffusion-limited transport and an exponential relationship indicating barrier-limited transport. We have observed, that proteins could take considerable time (sometimes as much as ~2 hours) to reach a dynamic equilibrium where the slope of the *C_R_* – *V* curves (i.e., referred to as sensitivity hereafter) would not change appreciably with each sweep (Figure 1e). Implications of non-equilibrated protein runs could severely distort the final output. For example, as seen in Figure 1e, the initial sweep resembles a scenario where events are missed at higher applied voltages due to fast protein translocations. However, further data acquisition suggests that this is merely due to the proteins not achieving the before-mentioned dynamic equilibrium within the nanopipette; possibly due to diffusion processes or electrostatic repulsion inside the taper. Thus, rather than relying on a single sweep, in this study, we performed multiple *C_R_* – *V* sweeps until a steady state was reached and the average of the steady-state condition was then treated as the final sensitivity for a given condition.

We then investigated the transport properties and structural implications across a broad range of pore sizes. The sensitivity with *d_p_* is shown in Figure 1g. We broadly categorized the pores into four zones—zone 1 (red region), zone 2 (blue region), zone 3 (yellow region), and, zone 4 (green region)—based on major breakpoints of sensitivity with *d_p_*. Zone 2 is more conventionally anticipated where sensitivity is seen to decrease with decreasing pore size due to the proportional relationship between capture radius and pore size.^22–23^ However, counterintuitively, an increase in sensitivity was seen in zone 1. This is thought to be due to voltage-mediated unfolding seen with proteins which would transform the proteins from their globular state to a more linear state. An intricate interplay of the hydrodynamic drag forces, structural properties, and electroosmotic flow profiles could be responsible for the observed pattern in this zone (i.e., zone 1). Pores in zone 1 produced the highest current drops. This was expected because the volume occlusion with respect to the size of the pore is highest in this region. The electrical signals from zone 1 were of sufficient SNR to be filtered using the highest available LPF setting of the Axopatch 200B (i.e., 100 kHz), and a representative trace corresponding to +300 mV-+1000 mV from a ~9 nm pore is shown in Figure 1h. The ability to reach higher LPF settings is important since the bandwidth governs the rise time of the filter and by extension whether a signal is attenuated or not. The rise time of the LPF is given by *T_r_* = 0.3321/*f_c_* where *f_c_* is the cutoff frequency of the LPF, with events faster than 2*T_r_* (i.e., translocation time < 2*T_r_*) been attenuated. For the more ubiquitous LPF setting (i.e., 10 kHz), 2*T_r_* is ~66 μs (the full-width half maximum (FWMH) approach allows to reach calculated durations as low as ~40 μs with reasonable errors).^16, 20, 24^ For the 100 kHz LP filter, the theoretical 2*T_r_* is ~6.6 μs (SI Figure S2). The translocation time distributions were then fitted with the first passage model: 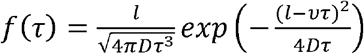 where *D, l*, and *ν* are the diffusion coefficient, effective sensing length, and drift velocity of the protein (see SI section 3 for more details). The peak of the fit (i.e., *τ_p_*)) corresponding to the events collected in Figure 1h fall above the attenuation limit of the 100 kHz LP filter (Figure 1i). However, upon filtering with the 10 kHz LPF, we see that the *τ_p_* for all voltages are within the attenuation zone of the filter as seen in Figure 1j. Thus, higher LPF (made possible by high SNR) enables the collection of nonattenuated events and thereby a non-distorted evaluation of protein characteristics.

As the pore size increase, the analyte confinement would decrease and by extension the probability of missing events. This would lead to a drop in sensitivity which is the case in zones 3 and 4. As the pore size is further increased (zone 4), we see a departure of *C_R_* – *V* relationship from the linear nature (Figure 2b). A nonlinear *C_R_* – *V* response is indicative of barrier-limited transport. However, the non-linearity in the *C_R_* – *V* response is not because of a reversal in the transport mechanism but due to poor SNR in larger pore sizes which is evident through the restoration of the diffusion-limited transport mechanism upon reducing the LP filtering to 5 kHz as seen in Figure 2c. To ensure that the *C_R_* – *V* pattern seen in Figure 2b is not due to undercounting of events, we varied the peak detection coefficient (PDC, the standard deviation of the baseline in the analysis window) from 5 (default value) to 4 in 0.5 steps. Decreasing the PDC below 4 could capture noise spikes from the baseline. As seen in Figure 2b, the shape of the *C_R_* – *V* patten is retained irrespective of the investigated PDC values which strengthen the notion that the observed deviation from the diffusion-limited transport is mainly due to poor SNR and not due to undercounting of events. Similarly, PDC was increased from the default value of 5 to 8 for the 5 kHz filtered trace as shown in Figure 2c. This was to see if the choice of PDC would impact the nature of the *C_R_* – *V* and as seen, with all the PDC values investigated, the linear relationship between *C_R_* and *V* was maintained. If the transport mechanism is truly barrier limited, the log of *C_R_* with *V* would be linear. However, as seen in Figure 2d, two distinct linear regions with a breakpoint can be seen. This is further evidence for the transport being not truly barrier limited and further emphasizes the need to scan a broad voltage range to identify the transport mechanism at play. It is thus vital to assess the influence of noise on the observed capture rate and identify applied voltages that are hampered by noise. One approach to calculating the noise floor is through the false event rate approach^8^ which was found to be ~0.03 events/s and ~0.3 events/s for the 10 kHz and 100 kHz LPFs. However, as shown in Figure S4a, this only classifies the lowest voltage as being hampered by the noise of the measurement. However, looking at the log of *C_R_* with *V* (Figure 2d), the second linear regime (low voltage) has a higher slope which could be arising due to false events. Thus, we defined the intersection point of the two linear regimes of the log *C_R_* – *V* curve as the ceiling of the noise floor (see SI section 4 for more details on the intersection point). Furthermore, the reversal of *C_R_* – *V* relationship from linear to non-linear is not limited to larger pores but is also seen with smaller pores filtered using a higher than optimal LPF as seen in Figure 2f. These observations further emphasize the underlying relationship between detectable events and the SNR and its implications on the transport mechanism (other examples are shown in Figure S5). Since the maximum available bandwidth ( ) is proportional to where and are voltage noise and total capacitance at the amplifier’s input and is correlated with the SNR, to operate at a higher bandwidth a higher (or lower) is needed.^25^ As such, when the SNR is sufficient, the predicted transport mechanism does not depend on the LPF setting as seen in Figure 2j and 2k.

**Figure 2:**
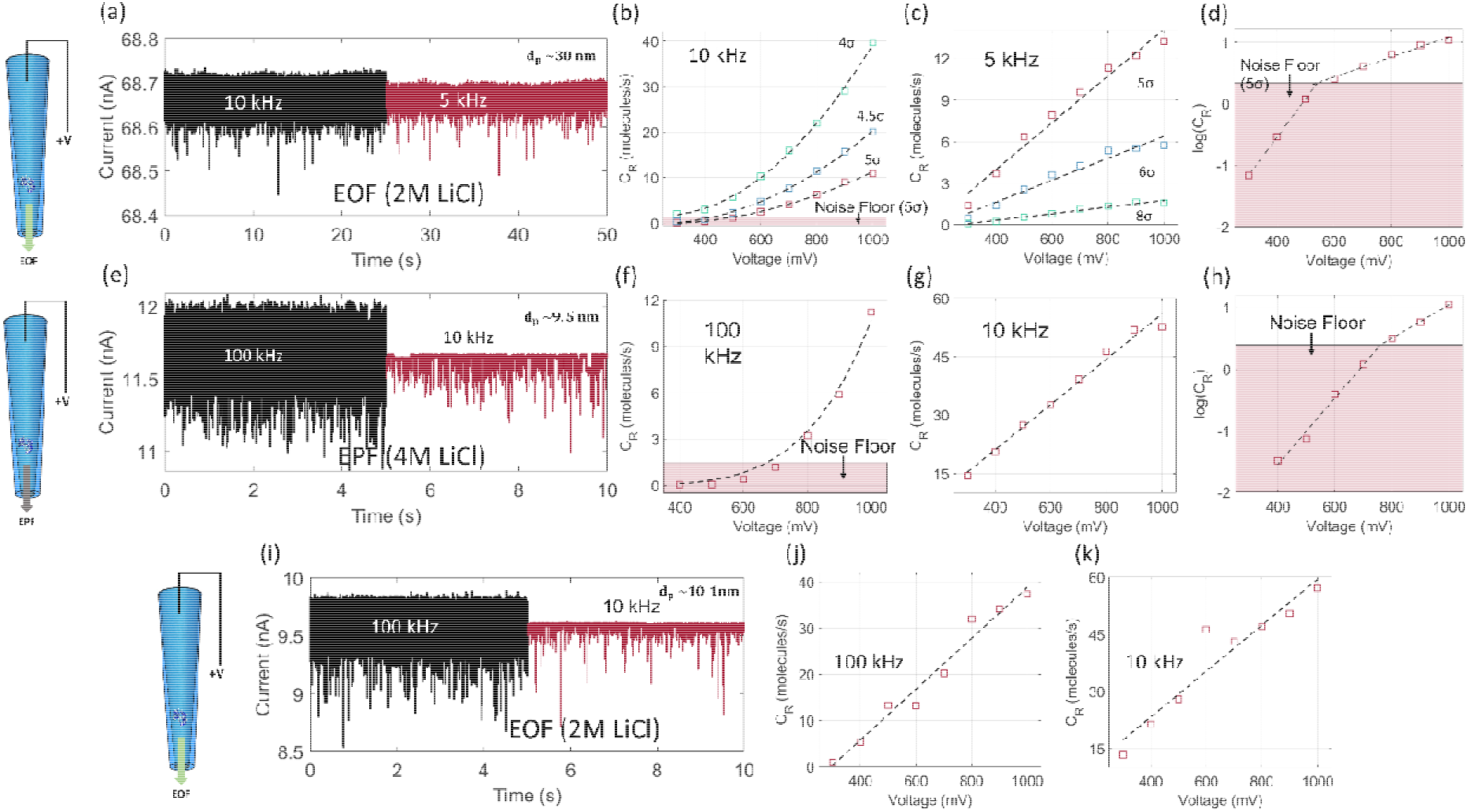
**(a)** Current trace (1000 mV) filtered at 10 kHz (black) and 5 kHz (marron), **(b)** with (10 kHz LPF), **(c)** with (5 kHz LPF) and **(d)** log of with V (10 kHz LPF) corresponding to ~30 nm pore. **(e)** Current trace (−1000 mV, presented in the positive bias resistive pulse direction for ease of comparison) filtered at 100 kHz (black) and 10 kHz (marron), **(f)** with (100 kHz LPF), **(g)** with at (10 kHz LPF) and **(h)** log of with (100 kHz LPF) corresponding to ~9.5 nm pore. **(i)** Current trace (1000 mV) filtered at 100 kHz (black) and 10 kHz (marron) and, with at **(j)** 100 kHz LPF, and **(k)** 10 kHz LPF corresponding to ~10.1 nm pore. The top row and bottom row experiments were done using 2M LiCl (electroosmotic transport) while those in the middle row were done using 4M LiCl (electrophoretic transport). All experiments were performed at pH ~8

### Protein Unfolding Kinetics

We limit the discussion hereafter to pore sizes excluding zone 4 due to insufficient SNR at 10 kHz LPF. Proteins are known to undergo voltage-induced unfolding and have been demonstrated widely using planar nanopores. ^17, 26–27^ The understanding of such (i.e., protein-unfolding) in nanopipettes is meager (relatively unexplored), and given the tapered geometry of nanopipettes, we sought to explore the behavior of proteins (with hSTf as the model protein) under a wide range of voltages and pore sizes spanning electroosmotic and electrophoretic dominant transport regimes. We chose hSTf as the model protein since it has been studied using silicon nitride nanopores in recent years. ^17, 26–27^ We would first discuss the electroosmotic dominant transport. All experiments in this regime were carried out using 2M LiCl. The protein was always placed in the pipette due to minimal analyte clogging (discussed later in the manuscript). We first looked at the ΔI distribution in zone 1 and zone 2. A broad ΔI distribution was observed (e.g., Figure S3b) which is typical for protein translocation, and they were fitted with a function in the form 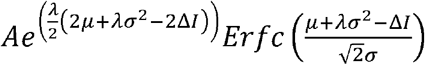 where *A, λ, μ σ*^2^ and *Erfc* are amplitude, rate of exponential component, means, variance, and complementary error function. The peak of this fit (*i.e*., ΔI_p_) showed an Ohmic relationship with the applied voltage for the pores in zone 2 (Figure 3a) which is indicative of the absence of voltage-driven unfolding. Similar behavior was seen for pores in zone 3 as well. The time constant (τ_0_) also showed a similar behavior with applied voltage as seen in Figure 3b. The time constant was computed by fitting the distributions shown in Figure S3d with a function in the form *A*_0_*e^-τ/τ_0_^* where *A*_0_ and *τ*_0_ are preexponential coefficient and time constant. Higher *τ*_0_ is indicative of a higher slower moving population and the decrease in *τ_0_* with increasing voltage reflects well on the increase in translocation speed with increasing voltage. The slope of the ΔI_p_ – *V* curves for zone 2 and zone 3 pores were plotted with pore diameter (*d_p_*, Figure 3c) which showed a weak linear relationship (R^2^ ~0.31). This is expected because if the protein is not unfolding, the ΔI_p_ would be dictated by the molecular volume of the native state (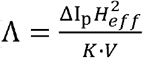 where Λ and, *H_eff_* molecular volume and effective length of the aperture)^28^ and ΔI_p_/V should be constant. Disregarding the outlier in Figure 3c (black boxed point), the mean value was found to be 0.024±0.006 pA/mV. Statistics further improved by setting an upper limit of 180 for the pull (*i.e*., disregarding the two points in the purple box of Figure 3c): 0.0026±0.004 pA/mV. Using this value, and assuming that the native state of hSTf is preserved under these conditions (molecular volume of hSTf ~144 nm^3^), *H_eff_* was found to be ~285±22 nm. This value was then used to calculate the pore size, *d_p_* (*L* = *H_eff_* in equation 1) and compared against the pore size evaluated from TEM imaging (*d*_*p*(*TEM*)_) where the two quantities for zone 1 pores were found to be within 10%.

**Figure 3:**
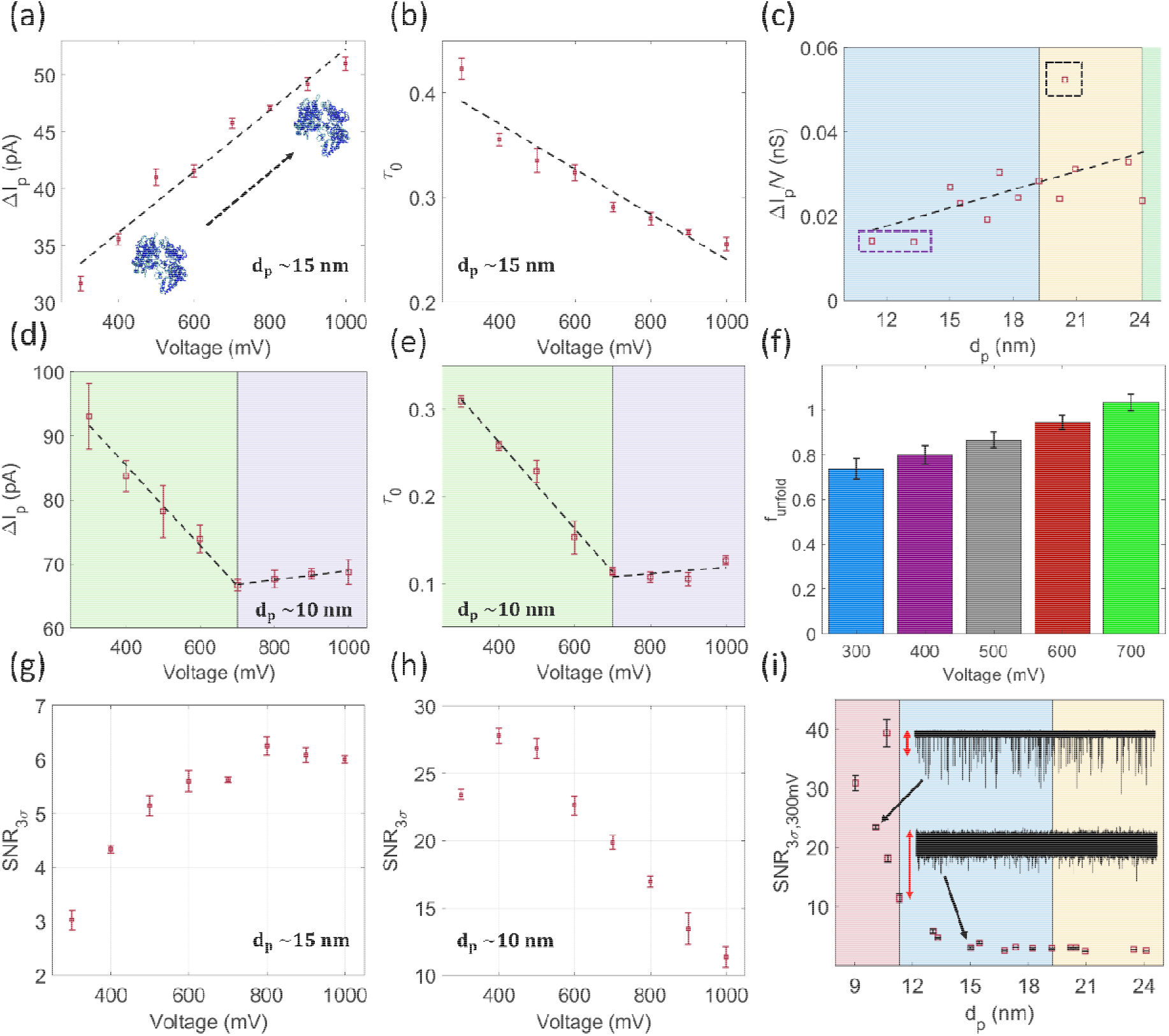
(**a**) Representative peak current drop ( ) and **(b)** time constant ( ) of a ~15 nm pore (zone 2). (**c**) Slope of vs applied voltage graphs corresponding to pores in zone 2 and 3. (**d**) Representative peak current drop ( ) and **(e)** time constant ( ) of a ~10 nm pore (zone 1). (**f**) The fraction of unfolding corresponding to 300, 400, 500, 600, and 700 mV from 10.1±0.8 nm diameter pores (4 independent trials). at 300 mV corresponding to (**g**) ~15 nm (zone 2), (**h**) ~10 nm (zone 1) diameter pores and, (**i**) all pores in zone 1-3. The vertical red arrows next to current traces in (i) represent 100 pA. All experiments were performed at pH ~8

Unlike pores in zones 2 and 3, the pores in zone 1 exhibited two distinct linear ranges: a decrease in ΔI_p_ (green region of Figure 3d) followed by a slight increase (purple region of Figure 3d). Since ΔI_p_ is related to Λ, a decrease in ΔI_p_ indicates a decrease in the Λ of the protein which is indicative of voltage-driven unfolding. The subsequent increase in ΔI_p_ could indicate that the protein has unfolded to a limit governed both by the protein and the conditions at play. Similar behavior was seen with τ_0_ as well (Figure 3e). After fitting the data points in each of the regions of Figure 3d with linear functions (shown as dashed lines), we defined the fraction of unfolding in the following manner 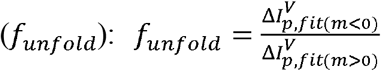 where 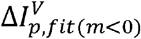 and 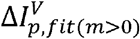 are ΔI_p_ at a given voltage computed from the positive and negative slopes. Thus, when the curve with the negative slope approaches the curve with the positive slope, *f_unfold_* would reach a value of 1. The *f_unfold_* for the pores in zone 1 are shown in Figure 3f and shows how the fraction of unfolded gradually increases with applied voltage (i.e., non-cooperative unfolding).

SNR can be defined as ΔI_p_/3σ_1_ where σ_1_ is the standard deviation of the open pore current.^25^ For uniformly charged linear analytes such as DNA, where the ΔI distribution is Gaussian-like, ΔI_p_ is more representative of SNR calculations. However, the ΔI distribution of proteins is broad and exponentially decays from the peak (Figure S3b). Thus, the ΔI_p_/ 3σ_1_ distribution was fitted with a Gaussian with an exponential tail and used the metric 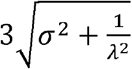 where 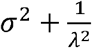 is the variance of the Gaussian with an exponential tail (Figure S3c) to define SNR as: 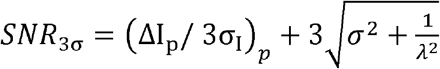 where (ΔI_p_/ 3σ_1_)_*p*_ is the peak of the ΔI_p_/ 3σ_1_ distribution. Thus, rather than SNR, we have defined a maximum SNR which encompasses the broad distribution of proteins. The *SNR*_3σ_ increased with increasing applied voltage for pores in zone 2 and 3 (Figure 3g). This is because the increase of ΔI_p_ with voltage outweighs that of σ_1_ thanks to the low dielectric noise properties of quartz. Moreover, this property allowed experiments to be conducted at higher voltages such as ±1000 mV without having to worry about open-pore noise which is not a luxury available with silicon nitride pores. *SNR*_3σ_ for pores in zone 1 (i.e., those that undergo voltage-drive unfolding) showed a decrease in *SNR*_3σ_ with voltage (Figure 3h). This is because, unlike pores in zone 2 and 3, pores in zone 1 induce protein unfolding which would lead to a decrease in ΔI_p_ with voltage as seen in Figure 3d. Thus, protein unfolding influences the behavior of ΔI_p_, *τ*_0_ and *SNR*_3σ_ and could be used as metrics to evaluate whether a protein is undergoing unfolding or not. The *SNR*_3σ_ at 300 mV is shown for all pores (Figure 3i) and we see that as the pore size decreases, *SNR*_3σ_ increases. This is anticipated since the percentage of volumetric occlusion by the analyte increase with decreasing pore size. This is also evident through the two traces shown in Figure 3i where the smaller pore showed deeper blockades compared to the larger pore.

### Transport Mode and Transport Direction

Next, we investigated the electrophoretic transport properties with zone 1 pores since voltage-driven unfolding and high *SNR*_3σ_ were present compared to other pore sizes. Interestingly, unlike in the EOF dominant cases, ΔI_p_ showed a linear relationship with applied voltage indicating that the protein is not unfolding under electrophoretic conditions (Figure 4b). In the EOF dominant realm, the EOF and EPF are in opposing directions whereas in the EPF dominant realm, due to the high salt concentration, the contribution of EOF is meager. Thus, the presence of the two opposing forces could facilitate the voltage-driven unfolding in the EOF dominant cases and the absence of such may allow the protein to translocate without any detectable unfolding. Furthermore, unlike in the EOF dominant realm, the sensitivity (i.e., the slope of *C_R_* – *V* curve) decreased with decreasing pore size (Figure 4d). This is more anticipated since the capture radius scales inversely with pore size. This further strengthens our hypothesis of the increase in sensitivity in zone 1 pores in the EOF dominant realm is influenced by a change in the protein conformation (i.e., voltage-driven unfolding). The inset of Figure 4d shows the sensitivity of EPF and EOF dominant transport conditions for comparable pores. As seen, in larger pores, EPF produces a higher sensitivity and eventually decreases below that produced by EOF dominant transport. This is attributed to an interplay between the capture radius and protein unfolding, which, as explained earlier, causes opposite trends in sensitivity for the two transport modes. *SNR_3σ300 mV_* (i.e., *SNR*_3σ_ at 300 mV), like with EO dominant cases, increased with decreasing pore size due to the increasing volumetric occlusion by the analyte (Figure 4e). However, as seen in Figure 4e, the *SNR*_3σ_ of EP dominant events (magenta squares) are less compared to their EO counterparts (black squares). A pure geometric argument taking volumetric occlusion into account would suggest an unfolded protein to produce a lower SNR compared to its globular counterpart. However, since Figure 4e involves two transport mechanisms, concluding solely based on volumetric occlusion and protein structure is challenging since the signal magnitude of proteins changes with electrolyte chemistry.^16^

**Figure 4:**
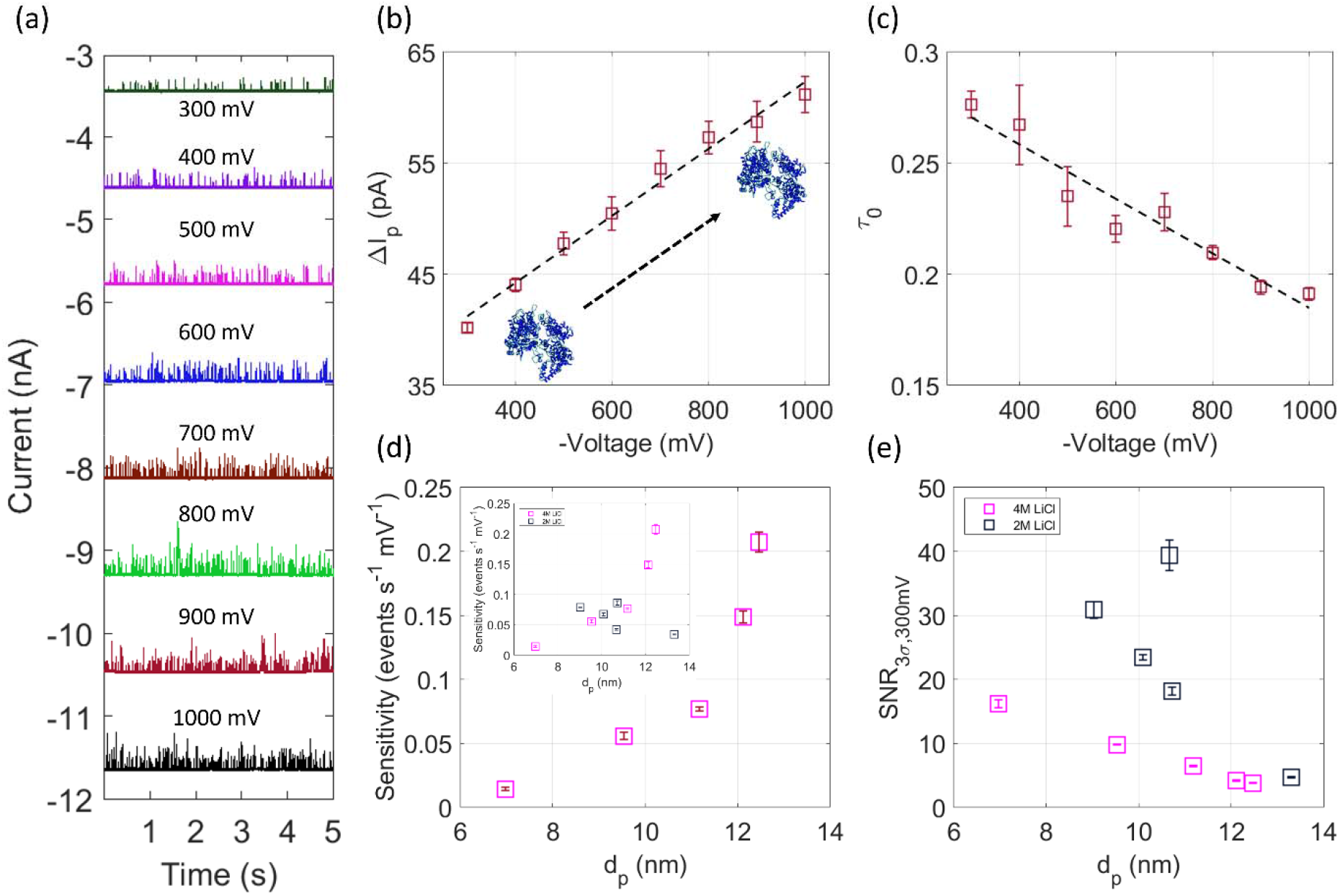
(**a**) 5-second representative current trace corresponding to hSTf translocating electrophoretically through a ~9.5 nm pore (forward direction, 4M LiCl, 10kHz LPF). (**b**) and (**c**) with applied voltage. (**d**) Sensitivity (i.e., the slope of vs applied voltage graph) and (**e**) with d_p_ corresponding to hSTf translocations in 4M LiCl (magenta) and 2M LiCl (black). The sensitivity corresponding to both 4M LiCl and 2M LiCl conditions is shown as an inset in (d). All experiments were performed at pH ~8

Up to now, the discussion was solely focused on the forward direction (i.e., pipette to bath translocations). However, nanopipettes have been shown to exhibit direction-dependent transport properties.^29^ Although such properties have been investigated with DNA, to the best of our knowledge, they have not been explored with proteins. Given the charge heterogeneity of proteins, voltage, pore size, electrolyte chemistry, and transport mechanism-dependent structural changes, it is important to explore the backward transport direction as well. A Representative current trace corresponding to hSTf translocating (in 4M LiCl) through a ~10.4 nm pore electrophoretically in the backward direction is shown in Figure 5a. The ΔI_p_ shows an Ohmic behavior with *V* indicating the absence of voltage-drive protein unfolding (Figure 5b). However, we have found the backward direction often leads to clogging (Figures 5d–5h). Although a considerable number of events are collectible through EPF dominant transport in the backward direction, clogging happens in EOF dominant cases before a significant amount of data can be collected (Table S1 and, Figures 5d–5e). To elucidate in the context of the number of events (using EOF transport of hSTf), a pore eventually clogged irreversibly (forward direction) produced >30,000 events while the best pore in the backward direction produced a mere ~4200 events.

**Figure 5:**
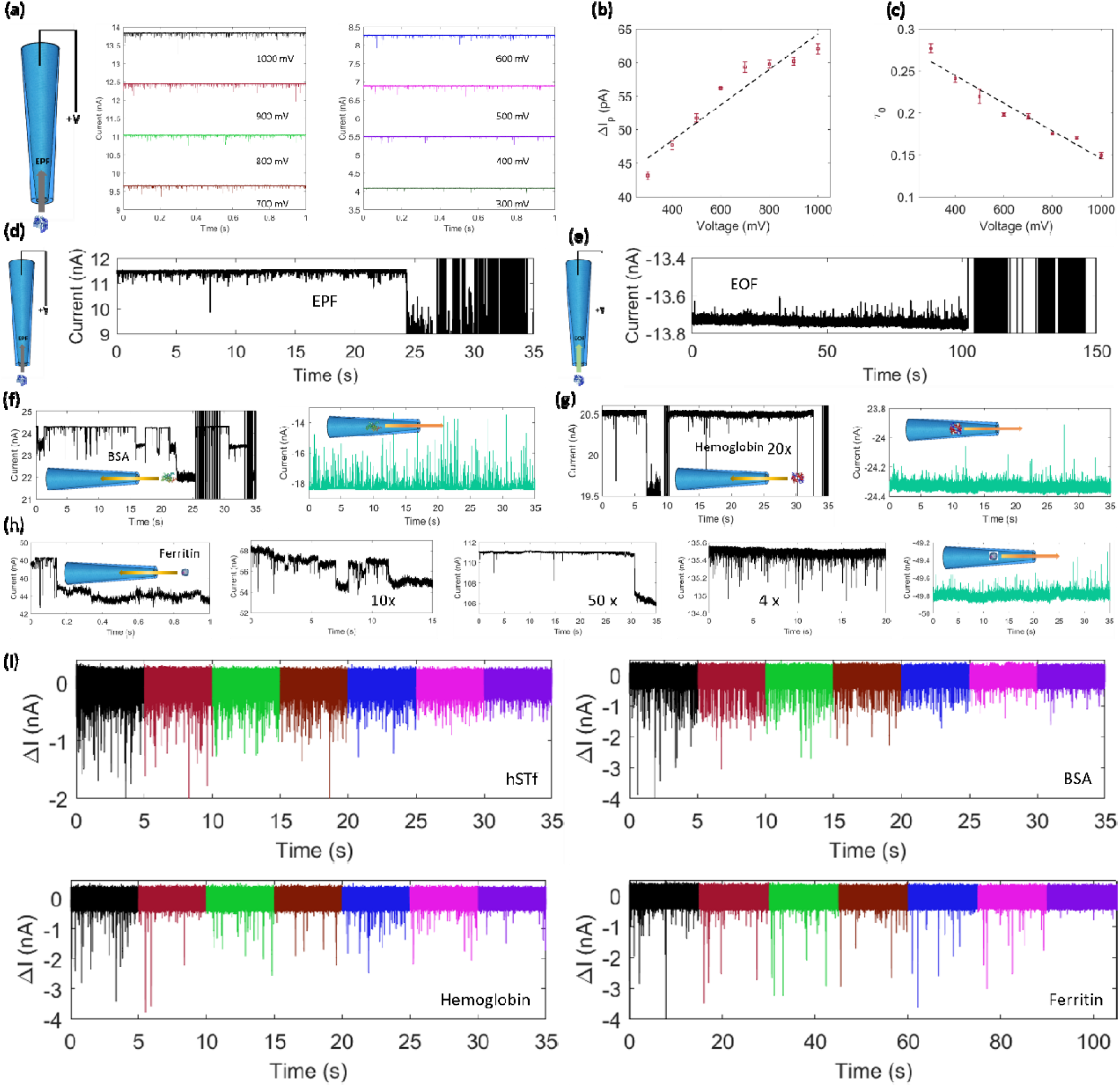
(**a)** Events from protein translocating electrophoretically in the backward direction (4M LiCl) through a ~10.4 nm pore with the corresponding **(b)** and **(c)** vs applied voltage. Example traces where frequent analyte clogging was observed for hSTf translocating in the backward direction through **(d)** electrophoresis (~11.3 nm pore) and **(e)** electroosmosis (~11.8 nm). Forward (black traces) and backward translocation (green traces) corresponding to **(f)** BSA (~20 nm, both pores), **(g)** hemoglobin (~21 and ~24 nm), and **(h)** ferritin (from left to right: ~19 nm, 27 nm, 32 nm, ~38 and, 22 nm). For hemoglobin and ferritin, the concentrations were diluted with respect to that used in the forward direction as noted in each figure. Table S1 outlines the total number of events observed in each of the cases in the figure. Experiments corresponding to hSTf and BSA were performed at pH ~8 while those of hemoglobin and ferritin were performed at pH ~10. (i) Translocations at 100 kHz LPF corresponding to (~9 nm pore), BSA (~9.2 nm pore), hemoglobin (~6.3 nm pore) and ferritin (~15.2 nm pore) in response to1000 mV (black), 900 mV (maroon), 800 mV (green), 700 mV (brown), 600 mV (blue), 500 mV (magenta) and 400 mV (purple).

Similar behavior in the backward direction was observed with BSA (Figure 5f), hemoglobin (Figure 5g), and ferritin (Figure 5h) as well. Of the four proteins, pore-clogging with ferritin was more severe where instantaneous irreversible clogging was observed (Figure 5h, only ~19 events were collectible). Upon diluting the ferritin by 10× and increasing the pore size from ~19 nm to ~27 nm, ~800 events could be collected before irreversibly clogging with the analyte. Further diluting ferritin (50× diluted) allowed a few more events to be collected (~1100 events using a ~32 nm pore). Increasing the pore size up to ~38 nm alleviated analyte clogging and permitted a higher ferritin concentration than the clogged cases to be used (4× diluted). An excess of 63,000 was collected and the pore remained open. The *C_R_* vs *V* for this larger pore resembled a barrier-limited scenario (Figure S5d) which was due to insufficient SNR (Figure S5e). A similar concentration-dependent clogging behavior was seen with hemoglobin as well where ~2,100 and ~3,800 were collected in the backward direction by diluting hemoglobin 5× and 20×, respectively. Analyte clogging could originate from a range of reasons such as co-translocations (*i.e*., two proteins entering at the same time), electrostatics (*e.g*., a positively charged protein adhering to a negative surface), pore-corking (*e.g*., pore diameter is smaller than the protein diameter). Since the size of the pores used for BSA (~7 nm), hemoglobin (~5 nm), and ferritin (~12 nm) are larger than the protein, pore corking could be ruled out. Although all the proteins are net-negatively charged, the charge heterogeneity of proteins could lead to adsorption through positively charged domains (i.e., electrostatics). Another avenue for clogging is co-translocations. Reducing the concertation of the protein could decrease the probability of co-translocations. Both co-translocations and electrostatics-mediated analyte-clogging is more prominent in smaller pores and become less severe as the size increase. Looking at the results presented in Figure 5, other than the geometry in the two transport directions, under similar conditions, the backward transport direction leads to more clogging. Moreover, events as many as ~600,000 have been collected with apo-ferritin from a single pipette in the forward direction (data not shown) and it is not uncommon to see protein translocations lasting for hours without any clogging issues. While in the backward directions, proteins can arrive at the pore in a multitude of directions or trajectories (i.e. the capture volume is nearly a full sphere rather than a hemisphere), while the tapered geometry may lead to some degree of alignment of the protein along the pore axis which could contribute to lowering co-translocations and by extension clogging. We speculate that the gradual confinement offered by the taper acts to promote single file ordering of the proteins. Thus, there could be fewer co-translocations in the forward direction compared to the backward direction which may lead to a longer pore lifetime. The ability and the know-how to refute pore clogging by the analyte and increase SNR were then used to record translocations at the highest available LPF setting of the Axopatch 200B (i.e., 100 kHz) for all four proteins used in this study (i.e., hSTf, BSA, hemoglobin, and ferritin). The low dielectric noise properties of quartz coupled with the ruleset we have formulated enabled the successful translocation of hSTf (~9 nm pore), BSA (~9.2 nm pore), hemoglobin (~6.3 nm pore), and ferritin (~15.2 nm pore) as shown in Figure 5i. The *τ_p_* (for 1000 mV) hSTf, BSA, hemoglobin, and ferritin were found to be ~13 μs, ~13 μs ~11 μs, and ~9 μs respectively which is slower than the theoretically predicted attenuation threshold of the 100 kHz LPF. We intend to expand this ruleset to a host of proteins spanning a wide range of sizes in a future study.

To gain a fundamental understanding of a few of the salient experimental observations seen in this study, numerical solving of Faraday’s electrostatic equations and the Navier-Stokes equations was performed. Briefly, an axisymmetric model was used and the inner diameters of the nanopipette were roughly based on those obtained by TEM. An insulating boundary is used for the nanopipette walls while ground and potential boundary conditions are used for the fluidic boundaries on either side of the nanopipette. Electroosmotic flow (EOF) was modeled using an electroosmotic boundary condition for the nanopipette walls where the critical input is the zeta potential. Importantly, the zeta potential of quartz varies with salt concentration and so the conductivity of the salt was incorporated into the model using this boundary condition. The electrostatics of the system, however, is not influenced by the conductivity of the electrolyte. For all the numerical models, a fixed value of electrophoretic mobility was used (1×10^-8^ m^2^/Vs) and the zeta potential was varied from −5 mV to −30 mV.

Most of the experiments in this study were performed using 2M LiCl which yielded EOF-driven events even despite the counteracting electrophoretic forces. EOF-driven events, especially under relatively high molar LiCl (e.g., >1M LiCl), have been reported elsewhere^16^ as well and seem to be the effect that disappears at higher salt concentrations such as 4M LiCl. Based on the Grahame equation (see Supporting Information), the zeta potential of the glass surface is most sensitive to electrolyte concentration at low salt (<1 M) and then undergoes a mild, and steady decline at higher salt concentrations. Since a relatively meager decrease in zeta potential is expected from 2M to 4M LiCl, we sought to explore the balance of EOF and EP-Drift within this range. The sensitivity of EOF-dominated transport on zeta potential was explored here and it was found that a zeta potential difference of less than 5 mV could alter the net transport of a protein. For the standard electrophoretic mobility used here, a zeta potential of −10 mV led to a peak EP drift velocity of approximately 70 mm/s and a peak EOF velocity of 50 mm/s yielding EP-dominance at the pore (**Figure 6a**). However, a zeta potential of −15 mV led to the same EP drift velocity and a higher EOF velocity of 80 mm/s leading to EOF-dominance. The sensitivity of EOF to zeta potential is clearly a critical factor in nanopore experiments, including those at high salt conditions such as 2M LiCl which was used in the present study.

**Figure 6:**
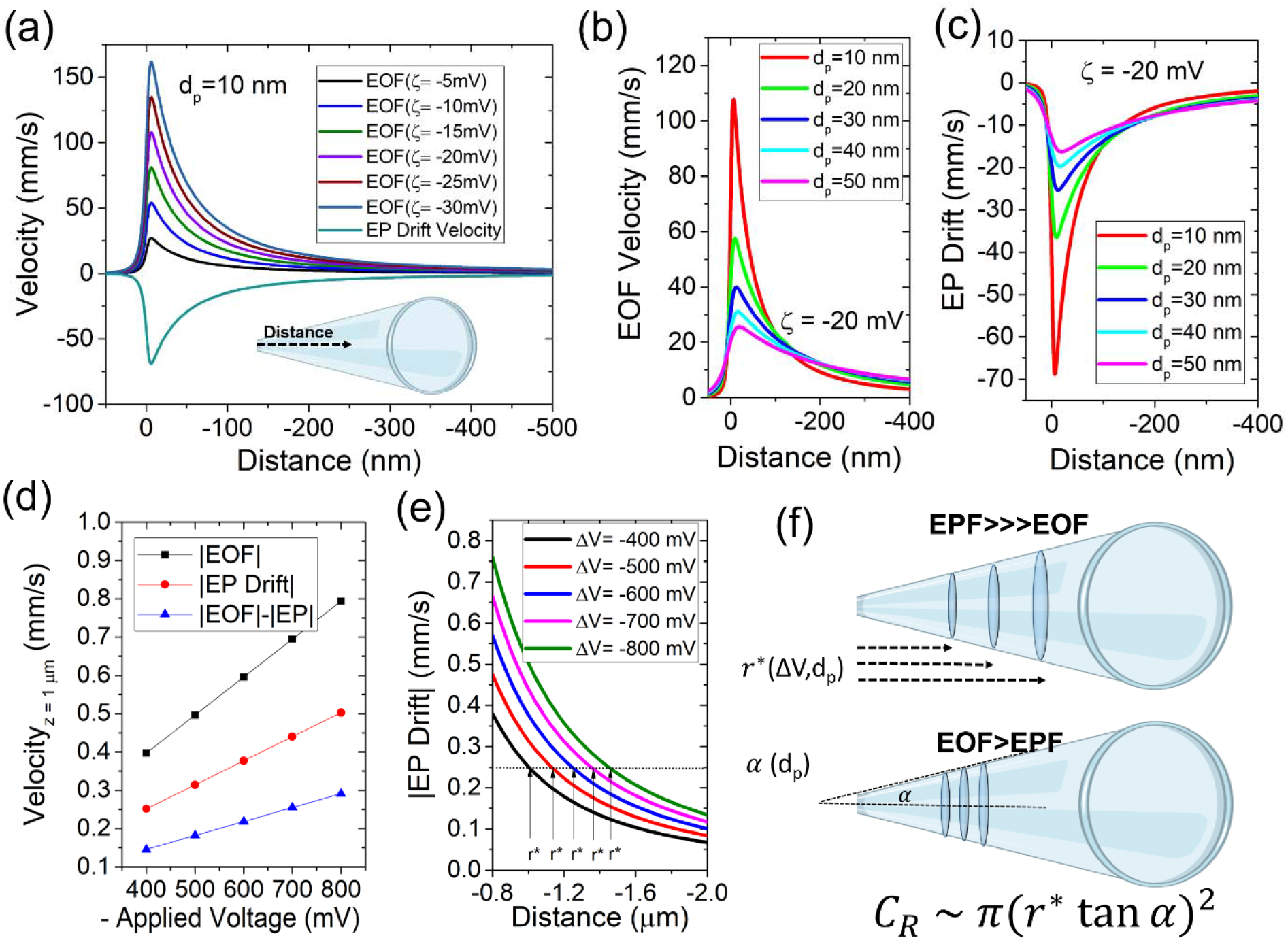
(a) Electroosmotic flow (EOF) velocities at different zeta potentials (−ζ = 5-30 mV) compared to the electrophoretic drift velocity of a protein with an electrophoretic mobility of 1×10^-8^ m^2^/Vs. The data was extracted from the axis of symmetry and labeled as the distance from the pore (inset). (b) EOF and (c) electrophoretic (EP) drift velocities plotted for different pore diameters (ζ = −20 mV for all simulations). (d) Net velocities of a hypothetical protein (1×10^-8^ m^2^/Vs) at a distance of 1 μm from the pore assuming EOF, EP drift, and both forces, respectively, and the varying levels of voltage responsiveness (i.e. sensitivity). (e) EP drift velocity of a protein is approximately 1 μm from the pore (d_p_=10 nm) at different voltage biases. (f) Schematic showing how capture radius (and capture volume) expands with voltage bias, pore diameter, and the half cone angle (α). An expression for capture rate is provided which is a geometric expression of the capture plane as it travels up the tapered region of the nanopipette.

Since EOF is also a function of the electric field inside the nanopipette, the EOF and EP drift velocities have nearly identical trends but are opposing one another for negatively charged analytes. This is due to the cations (Li^+^) being positive whereas the analyte is negatively charged. The convention used here was that EOF is positive and EP drift velocity is negative to show that they are opposing one another (**Figure 6b**). As the pore size is increased, both EOF and EP drift velocity decreases inside the pore but increases far away (>200nm) from the pore. The cross-over point is approximately located 150 nm from the nanopipette tip. Therefore, for both EOF and EP-dominant transport, the capture radius increases as the pore increases in diameter and by extension, *C_R_* should also, therefore, increase with *d_p_*. Since EP drift and EOF counteract each other, the relative net analyte motion (EP+EOF) will directly influence the sensitivity to voltage (a metric that was quantified in the experimental part of this study). Recalling that sensitivity is the slope of the *C_R_* versus voltage curve, an analogous relationship is observed considering EP drift, EOF, and net migration at a specified distance (1μm) from the pore (**Figure 6c**). The true capture radius (r*) is found using a certain threshold for the migration speed that can overcome diffusion. For illustration purposes, an arbitrary threshold of 0.25 m/s was set for the purely EP-dominant capture scenario (**Figure 6c**).

It is clear that capture radius increases into the nanopipette tapered region as voltage increases and *d_p_* increases however the exact relationship (e.g., linear, exponential, etc.) has not been investigated. As the capture radius extends up the tapered region of the nanopipette, the hemispherical surface area increases non-linearly. Assuming that the hemisphere can be modeled as a flat planar surface, the surface area of a circular plane will extend and rapidly grow as the capture radius increases. The relationship can therefore be written as *C_R_* ≈ *π*(*r\** tan *α*)^2^ where α is the half cone angle. Since α was observed to increase with increasing pore diameter (**Figure 1b**), α is also a critical factor that can increase the *C_R_* and therefore the sensitivity parameter calculated in our experiments. The factors that seem to increase sensitivity include the *d_p_* and α whereas conditions where EOF and EP drift are counteracting, the sensitivity would be reduced (**Figure 6d**). This roughly coincides with **Figure 4d** wherein EP-dominant transport showed a clear exponential increase in sensitivity for the d_p_~7-12.5 nm pore size range whereas EOF-dominant transport was lacking an exponential increase. The location of capture and the dynamics with respect to the applied voltage is observed to be non-linear within the nanopipette system and further investigation into the sensitivity parameter could help understand the relative contributions of EP drift and EOF in the future.

## Concluding Remarks

In this study, we have discussed a detailed framework to refute pore clogging by the analyte and improve SNR by focusing on the influence of pore size, electric field, transport direction (i.e., forward, and backward), and mechanism (electroosmosis and electrophoresis) both on the translocation process and the conformation of proteins (i.e., folded vs unfolded). Pore clogging by the analyte is a legacy issue with nanopores which negatively impacts the lifetime of a pore and its ability to collect large amounts. While both are crucial in their venture to translate into the commercial space the latter is critical for approaches such as machine learning to gain more insight into event signatures. In this study, absent any surface functionalization methods, we have developed a framework to achieve two core goals: refuting pore clogging and increasing SNR. To achieve these two goals, a host of factors were tuned and optimized: electrolyte chemistry, pore size, and transport direction. These optimization parameters directly influenced the transport mechanism (2M LiCl showed electroosmotic events while 4M LiCl showed electrophoretic events), SNR (smaller pores showed the highest SNR and with such pores, electroosmotic events showed a higher SNR), sticking probability (forward transport direction was reliant against clogging while the backward transport direction was more prone to clogging). Thus, the above observations could be condensed into,

i. Forward transport direction is more resilient against pore-clogging while backward transport direction is prone to analyte clogging. The ability to access the pore from a multitude of directions promoting co-translocations was thought to be the reason behind enhanced clogging probability in the backward direction while the tapered geometry could be hydrodynamically aligning the analytes with the pore axis and by extension lowering the clogging probability in the forward direction.
ii. With forward transport direction, smaller pore sizes can be explored to improve SNR to enable high bandwidth data acquisition (100 kHz LPF with the Axopatch 200B) to alleviate temporal signal attenuation. The pore size spanned well into small pores that are <2× the size of proteins where sufficient confinement and SNR led to high bandwidth studies in the future.
iii. (ii) is also facilitated by the low dielectric noise properties of quartz. Otherwise, with increasing LPF, the increase in open-pore noise could overwhelm the signal amplitude.
iv. (iii) and (i) enable the data acquisition at higher voltages which otherwise (more often) lead to intermittent and thereafter irreversible analyte clogging (prevented by (i)) and unstable open-pore currents (prevented by (iii)).
v. The SNR increase in (ii) is also governed by the electrolyte chemistry where for hSTf, 2M LiCl was seen to showcase a higher SNR compared to 4M LiCl which was attributed to the difference in the transport mechanism associated with the two salts.

This optimization ruleset enabled to record >20,000 events/pipette very conveniently and often allowed the acquisition of >50,000 events/pipette. We have even demonstrated >100,000 events/pipette which if let to run longer could even achieve >500,000 events/pipette as was seen with apo-ferritin. To the best of our knowledge, such statistics are not documented with nanopipettes and would pave the way for these sensors to venture into the commercial space. While exploring the optimal conditions to achieve such impressive event statistics, a few more parameters that are often overlooked came under scrutiny:

a. Voltage must be swept until the slope of *C_R_* – *V* (i.e., sensitivity) does not change appreciably with each voltage sweep.
b. Appropriate LPF must be used to process the data. A higher than optimal LPF could lead to distortion of *C_R_* – *V* curves mainly due to missing events due to poor SNR.

Both (a) and (b) are often overlooked in literature perhaps due to their ostensible triviality. However, we see that (a) changes over time and sometimes takes as much as ~2 hrs to reach a steady value and the sensitivity of this steady state is often a few fold higher than the initially recorded sensitivity value. On the other hand, overlooking (b) could lead to misinterpretation of the energetics of the transport mechanism (i.e., diffusion vs barrier limited). With appropriate LPF, we found that the transport through all pore sizes was diffusion limited and without such optimization, especially with larger pores, the transport seemed to be barrier limited.

The framework for optimizing translocation enabled the study of the implication of the external parameters on the structure of the core morel protein, hSTf. Voltage-induced unfolding was seen with zone 1 pores (with EO dominant events) whereas the Δ*I_p_* – *V* relationship was Ohmic in other zones indicating the absence of such unfolding. On the contrary, no voltage-driven unfolding was seen with EP dominant events even with zone 1 pores. In the EO realm, the EO and EP forces are opposing whereas in the EP dominant realm, due to the high salt (i.e., 4M LiCl), the opposing EO is meager. Thus, our findings showcased a framework for optimal translocation conditions (e.g., resilience against pore clogging and high SNR for high bandwidth recordings) for proteins and the implications of the optimization parameters on the translocation properties and the protein structure. We believe these findings could help improve future nanopore studies and overcome legacy issues associated with the technology.

## Acknowledgments

This work was partially supported by the Human Frontier Science Program (RGY0066/2018). The authors would like to thank the University of California at Riverside for the software suites provided for this study.

## Author Contributions

KJF formulated the idea and, YMNDYB carried out the experiments and data analysis. KJF performed the simulation. YMNDYB and KJF wrote the manuscript.

## Ethics Declarations

The authors declare no competing interests.

## Notes

### Competing Interest Statement

The authors have declared no competing interest.

## References

1. Howorka, S.; Siwy, Z. S., Reading amino acids in a nanopore. Nature Biotechnology 2020, 38 (2), 159–160.

2. Goto, Y.; Akahori, R.; Yanagi, I.; Takeda, K.-i., Solid-state nanopores towards single-molecule DNA sequencing. Journal of Human Genetics 2020, 65 (1), 69–77.

3. Karawdeniya, B. I.; Bandara, Y. M. N. D. Y.; Khan, A. I.; Chen, W. T.; Vu, H.-A.; Morshed, A.; Suh, J.; Dutta, P.; Kim, M. J., Adeno-associated virus characterization for cargo discrimination through nanopore responsiveness. Nanoscale 2020.

4. McMullen, A.; De Haan, H. W.; Tang, J. X.; Stein, D., Stiff filamentous virus translocations through solid-state nanopores. Nature Communications 2014, 5, 4171.

5. Karawdeniya, B. I.; Bandara, Y. N. D.; Nichols, J. W.; Chevalier, R. B.; Dwyer, J. R., Surveying silicon nitride nanopores for glycomics and heparin quality assurance. Nature Communications 2018, 9 (1), 3278.

6. Chuah, K.; Wu, Y.; Vivekchand, S.; Gaus, K.; Reece, P. J.; Micolich, A. P.; Gooding, J. J., Nanopore blockade sensors for ultrasensitive detection of proteins in complex biological samples. Nature Communications 2019, 10 (1), 1–9.

7. Zhu, Z.; Duan, X.; Li, Q.; Wu, R.; Wang, Y.; Li, B., Low-noise nanopore enables in-situ and label-free tracking of a trigger-induced DNA molecular machine at the single-molecular level. Journal of the American Chemical Society 2020, 142 (9), 4481–4492.

8. Plesa, C.; Kowalczyk, S. W.; Zinsmeester, R.; Grosberg, A. Y.; Rabin, Y.; Dekker, C., Fast translocation of proteins through solid state nanopores. Nano Letters 2013, 13 (2), 658–63.

9. Rosenstein, J. K.; Wanunu, M.; Merchant, C. A.; Drndic, M.; Shepard, K. L., Integrated nanopore sensing platform with sub-microsecond temporal resolution. Nature Methods 2012, 9 (5), 487.

10. Li, J.; Hu, R.; Li, X.; Tong, X.; Yu, D.; Zhao, Q., Tiny protein detection using pressure through solid-state nanopores. Electrophoresis 2017, 38 (8), 1130–1138.

11. Yusko, E. C.; Johnson, J. M.; Majd, S.; Prangkio, P.; Rollings, R. C.; Li, J.; Yang, J.; Mayer, M., Controlling protein translocation through nanopores with bio-inspired fluid walls. Nature Nanotechnolagy 2011, 6 (4), 253–260.

12. Tabard-Cossa, V.; Trivedi, D.; Wiggin, M.; Jetha, N. N.; Marziali, A., Noise analysis and reduction in solid-state nanopores. Nanotechnology 2007, 18 (30), 305505.

13. Chen, P.; Mitsui, T.; Farmer, D. B.; Golovchenko, J.; Gordon, R. G.; Branton, D., Atomic layer deposition to fine-tune the surface properties and diameters of fabricated nanopores. Nano Letters 2004, 4 (7), 1333–1337.

14. Lee, M.-H.; Kumar, A.; Park, K.-B.; Cho, S.-Y.; Kim, H.-M.; Lim, M.-C.; Kim, Y.-R.; Kim, K.-B., A low-noise solid-state nanopore platform based on a highly insulating substrate. Scientific Reports 2014, 4, 7448.

15. Welch, S., Transferrin: the iron carrier. CRC Press: 1992.

16. Saharia, J.; Bandara, Y. N. D.; Karawdeniya, B. I.; Hammond, C.; Alexandrakis, G.; Kim, M. J., Modulation of electrophoresis, electroosmosis and diffusion for electrical transport of proteins through a solid-state nanopore. RSC Advances 2021, 11 (39), 24398–24409.

17. Saharia, J.; Bandara, Y. N. D.; Goyal, G.; Lee, J. S.; Karawdeniya, B. I.; Kim, M. J., Molecular-Level Profiling of Human Serum Transferrin Protein through Assessment of Nanopore-Based Electrical and Chemical Responsiveness. ACS Nano 2019, 13 (4), 4246–4254.

18. Saharia, J.; Bandara, Y. N. D.; Kim, M. J., Investigating protein translocation in the presence of an electrolyte concentration gradient across a solid□state nanopore. Electrophoresis 2022.

19. Bandara, Y. N. D.; Farajpour, N.; Freedman, K. J., Nanopore Current Enhancements Lack Protein Charge Dependence and Elucidate Maximum Unfolding at Protein’s Isoelectric Point. Journal of the American Chemical Society 2022, 144 (7), 3063–3073.

20. Bandara, Y. N. D.; Saharia, J.; Karawdeniya, B. I.; Kluth, P.; Kim, M. J., Nanopore Data Analysis: Baseline Construction and Abrupt Change-Based Multilevel Fitting. Analytical Chemistry 2021, 93 (34), 11710–11718.

21. Sun, L.; Shigyou, K.; Ando, T.; Watanabe, S., Thermally driven approach to fill sub-10-nm pipettes with batch production. Analytical Chemistry 2019, 91 (21), 14080–14084.

22. Wanunu, M.; Morrison, W.; Rabin, Y.; Grosberg, A. Y.; Meller, A., Electrostatic focusing of unlabelled DNA into nanoscale pores using a salt gradient. Nature nanotechnology 2010, 5 (2), 160–165.

23. Rowghanian, P.; Grosberg, A. Y., Electrophoretic capture of a DNA chain into a nanopore. Physical Review E 2013, 87 (4), 042722.

24. Pedone, D.; Firnkes, M.; Rant, U., Data analysis of translocation events in nanopore experiments. Analytical Chemistry 2009, 81 (23), 9689–9694.

25. Shekar, S.; Chien, C.-C.; Hartel, A.; Ong, P.; Clarke, O. B.; Marks, A.; Drndic, M.; Shepard, K. L., Wavelet denoising of high-bandwidth nanopore and ion-channel signals. Nano Letters 2019, 19 (2), 1090–1097.

26. Freedman, K. J.; Haq, S. R.; Edel, J. B.; Jemth, P.; Kim, M. J., Single molecule unfolding and stretching of protein domains inside a solid-state nanopore by electric field. Scientific Reports 2013, 3, 1638.

27. Freedman, K. J.; Ju□rgens, M.; Prabhu, A.; Ahn, C. W.; Jemth, P.; Edel, J. B.; Kim, M. J., Chemical, thermal, and electric field induced unfolding of single protein molecules studied using nanopores. Analytical Chemistry 2011, 83 (13), 5137–5144.

28. Sharma, V.; Freedman, K. J., Pressure-Biased Nanopores for Excluded Volume Metrology, Lipid Biomechanics, and Cell-Adhesion Rupturing. ACS Nano 2021.

29. Chen, K.; Bell, N. A.; Kong, J.; Tian, Y.; Keyser, U. F., Direction-and salt-dependent ionic current signatures for DNA sensing with asymmetric nanopores. Biophysical Journal 2017, 112 (4), 674–682.

