## Supporting Information for "Over Two Million Protein Translocations Reveal Optimal Conditions for High Bandwidth Recordings, Studying Transport Energetics, and Unfolding"

### Section 1: Size Calibration of 0.5 mm ID Quartz Pipettes

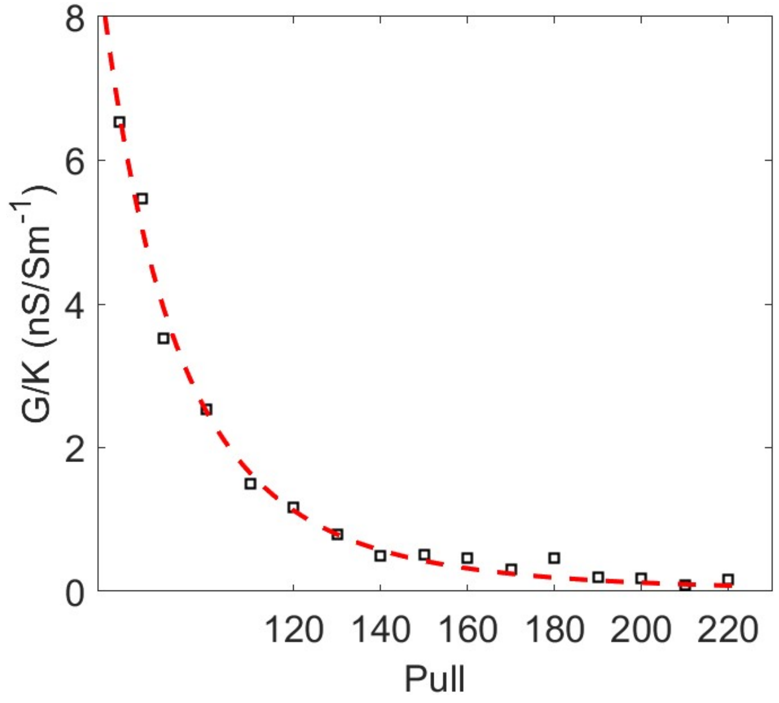

**Figure S1:** Pull vs G/K for pores fabricated using 0.5 mm internal diameter quartz pipettes. Raw data (hollow squares) were fitted with a function in the form $P = (G/K)^n$ where $P$ and $n$ are pull and an arbitrary exponent for fitting purposes. $n$ was found to be ~-4.4.

### Section 2: Calibration of the 100 kHz lowpass filter setting

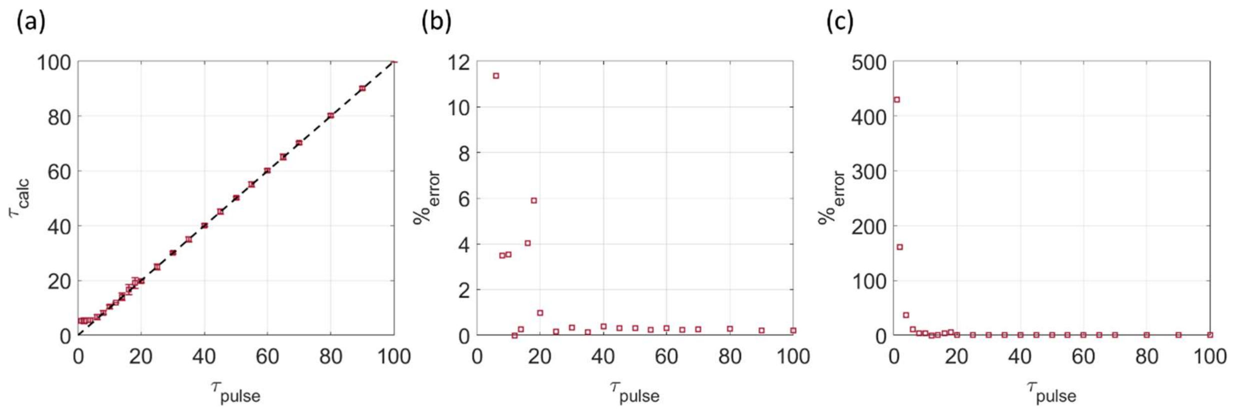

**Figure S2:** (a) Calibrating the 100 kHz lowpass filter of the Axopatch 200B using a function generator with $\tau_{pulse}$ and $\tau_{calc}$ been pulse width fed by the function generator and that computed by the analysis platform respectively. The dashed line corresponds to the ideal fit (slope =1). Percentage error ($\%_{error}$) from **(b)** $\tau_{pulse}$ = 6 μs onwards and **(c)** $\tau_{pulse}$ = 1 μs.

Pulses were fed to the rear switch of the Axopatch 200B using a function generator (Siglent SDG 1032x). The lowpass filter was set at 100 kHz. Data were sampled using the maximum available frequency of Digidata 1550B (i.e., 500 kHz) and analyzed using the *EventPro* platform which uses the FWHM (full width at half maximum) to compute the pulse width. Pulses as low as 1μs were used. While the percentage error ($\%_{error}$) was at a reasonable <10% down to 8 μs, it (i.e., $\%_{error}$) was slightly above 10% at 6 μs and exponentially thereafter as seen in Figure S2c. Thus, the lowest detectable pulse duration with reasonable deviation from the expected pulse width is ~6 μs, which is in good agreement with the value computed for the 100 kHz LPF (i.e., ~6.6 μs).

### Section 3: Model Fittings

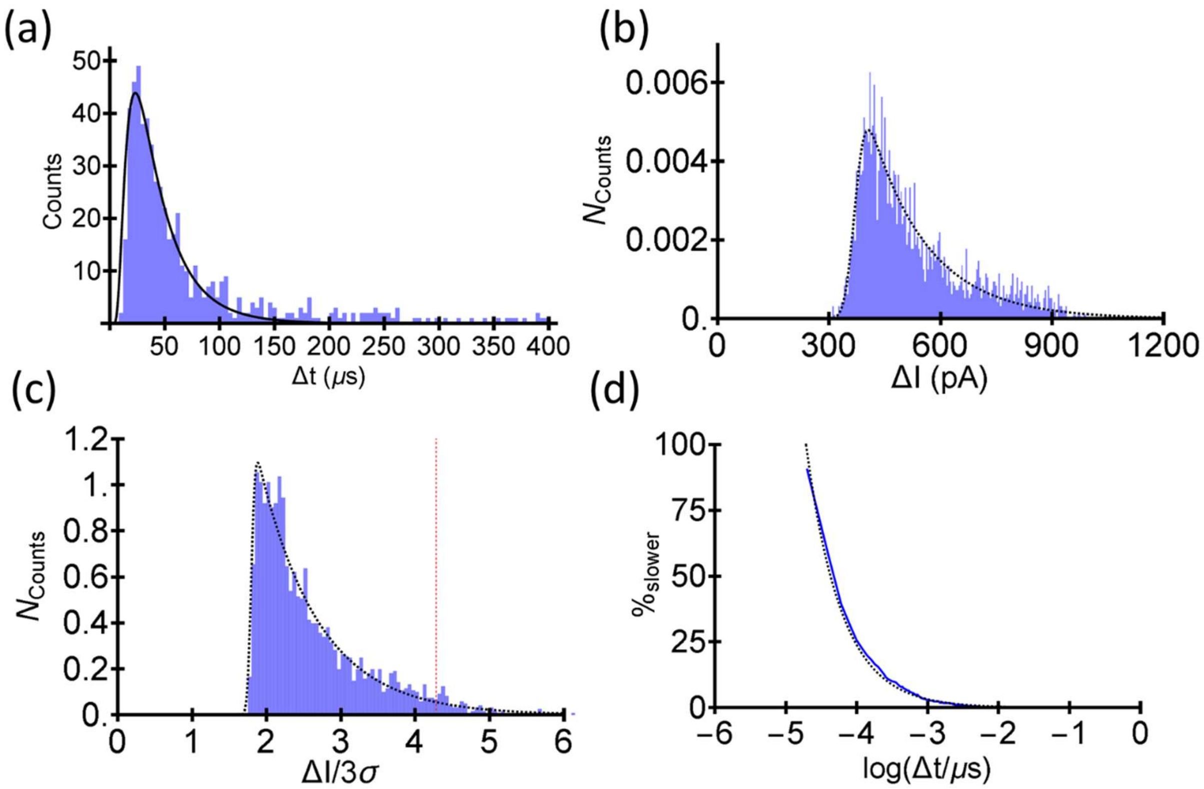

**Figure S3:** Example histograms/plots corresponding to **(a)** translocation time (Δt), (**b)** current drop (ΔI) **(c)** signal-to-noise ratio (ΔI/3σ) and **(d)** % slower events. Black lines correspond to the fits. The Red dashed line in (c) corresponds to the $SNR_{max}$

All histograms and their corresponding fits were constructed using Mathematica. For fitting the inbuilt *Non-linear-Model-Fit* function was used.

% slower events: The percentage of events that are slower than the value defined by the x-axis.

### Section 4: Noise Floor

The noise floor can be calculated using the following,

$C_{R,false} = kf_c e^{\left(-\frac{PDC^2}{2I_{rms}^2}\right)}$ where $C_{R,false}$, $k$, $f_c$, $PDC$ and $I_{rms}$ are false event rate, a constant (0.849), bandwidth, peak detection coefficient, and rms current.

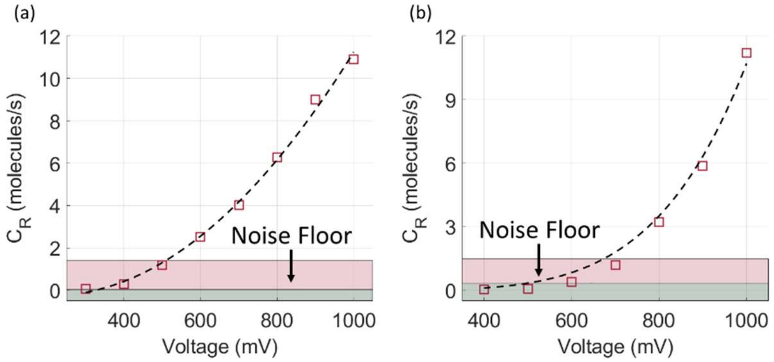

**Figure S4:** Capture rate with voltage for the pores shown in **(a)** Figure 2a (10 kHz LPF and PDC of 5) and **(b)** Figure 2e (100 kHz LPF and PDC of 5) of the main text. The green and red are noise floors calculated using the false capture rate approach and the intersection approach discussed in the main text.

**Identifying the Intersection point:** To identify the breakpoints in Figures 2d, 2h S5c, and S5f, the *findchangepts* function of MATLAB (with linear statistics) which identifies abrupt changes in a set of data points using the statistics specified. In these cases, linear statistics were used which would identify slope-based changes. The maximum number of change points was set to 1 since there is only one breakpoint.

Other examples where a reversal of $C_R - V$ is observed due to higher than optimal LPF:

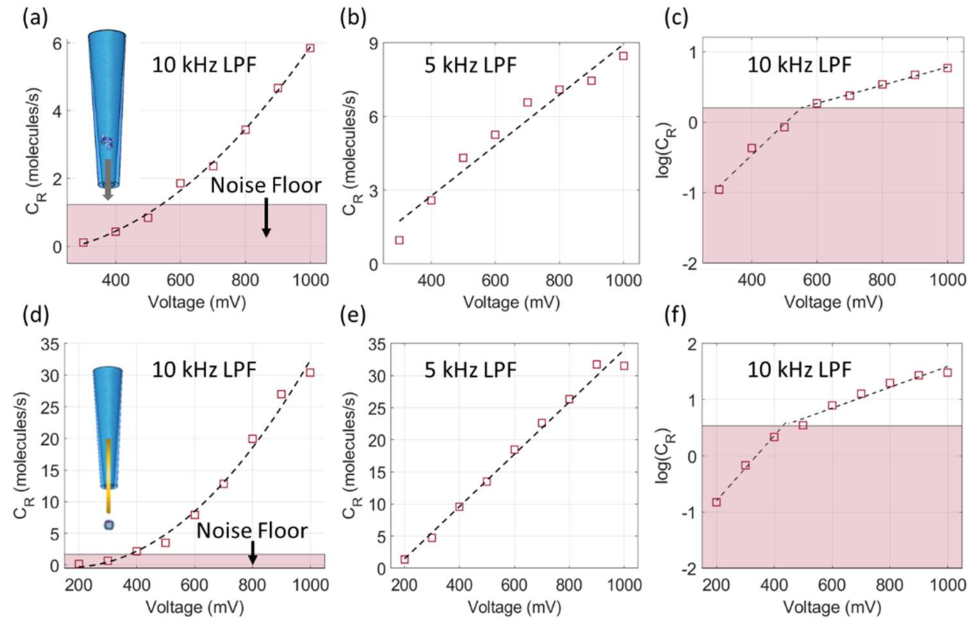

**Figure S5:** **(a)** $C_R$ with $V$ (10 kHz LPF), **(b)** $C_R$ with $-V$ (5 kHz LPF) and **(c)** log of $C_R$ with V (10 kHz LPF) corresponding to hSTf translocations through a ~26.7 nm pore in the forward direction (electroosmosis dominant). **(d)** $C_R$ with $V$ (10 kHz LPF), **(e)** $C_R$ with $V$ (5 kHz LPF) and **(f)** log of $C_R$ with V (10 kHz LPF) corresponding to ferritin translocations through a ~37.5 nm pore in the backward direction (electrophoresis dominant).

### Section 5: Event Log

| Protein | ~[Protein] (nM) | [LiCl] (M) | Transport Direction | G (nS) | Events | Comments |
|---|---|---|---|---|---|---|
| hSTf | 600 | 2 | Forward | 55.6 | 19,806 | Open |
| hSTf | 600 | 2 | Forward | 68.4 | 32,618 | Open |
| hSTf | 600 | 2 | Forward | 47 | 36,701 | Open |
| hSTf | 600 | 2 | Forward | 25.7 | 136,866 | Open |
| hSTf | 600 | 2 | Forward | 19.6 | 79,038 | Open |
| hSTf | 600 | 2 | Forward | 11.5 | 27,773 | Open |
| hSTf | 600 | 2 | Forward | 24.1 | 68,694 | Open |
| hSTf | 600 | 2 | Forward | 15.1 | 63,028 | Open |
| hSTf | 600 | 2 | Forward | 44.7 | 46,417 | Open |
| hSTf | 600 | 2 | Forward | 36.3 | 33,542 | Clogged |
| hSTf | 600 | 2 | Forward | 34.8 | 67,024 | Open |
| hSTf | 600 | 2 | Forward | 31.1 | 103,342 | Open |
| hSTf | 600 | 2 | Forward | 28.1 | 93,594 | Open |
| hSTf | 600 | 2 | Forward | 20.7 | 92,568 | Open |
| hSTf | 600 | 2 | Forward | 33.9 | 58,447 | Open |
| hSTf | 600 | 2 | Forward | 10.3 | 73,729 | Open |
| hSTf | 600 | 2 | Forward | 15.6 | 62,274 | Open |
| hSTf | 600 | 2 | Forward | 10.4 | 83,778 | Open |
| hSTf | 600 | 2 | Forward | 9.2 | 109,374 | Open |
| hSTf | 600 | 2 | Forward | 7.5 | 98,653 | Became Unstable |
| hSTf | 600 | 4 | Forward | 19.9 | 304,443 | Open |
| hSTf | 600 | 4 | Forward | 16.2 | 172,773 | Open |
| hSTf | 600 | 4 | Forward | 6.5 | 263,19 | Became bit sticky |
| hSTf | 600 | 4 | Forward | 18.9 | 96,935 | Open |
| hSTf | 600 | 4 | Forward | 11.9 | 83,284 | Became bit sticky |
| hSTf | 600 | 2 | Backward | 12.4 | 570 | Permanently Clogged |
| hSTf | 600 | 2 | Backward | 6.7 | 4,267 | Frequent Clogging |
| hSTf | 600 | 4 | Backward | 14.0 | 130,329 | Open |
| hSTf | 600 | 4 | Backward | 16.4 | 21,549 | Clogged |
| hSTf | 600 | 4 | Backward | 15.0 | 125,594 | Open |
| hSTf | 600 | 4 | Backward | 12.7 | 66,342 | Unstable |
| BSA | 600 | 4 | Backward | 49 | 47,051 | Permanently Clogged |
| BSA | 600 | 4 | Forward | 46.7 | 80,596 | Open |
| Hemoglobin (1×) | 3100 | 4 | Forward | 62.1 | 139,561 | Open |
| Hemoglobin (5×) | 620 | 4 | Backward | 49.8 | 2,157 | Permanently Clogged |
| Hemoglobin (20×) | 125 | 4 | Backward | 50.9 | 3,839 | Very Unstable |
| Ferritin (1×) | 68 | 4 | Forward | 49.9 | 119,843 | Open (clogs occasionally) |
| Ferritin (1×) | 68 | 4 | Backward | 47.9 | 19 | Permanently Clogged |
| Ferritin (10×) | 7 | 4 | Backward | 88.6 | 801 | Permanently Clogged |
| Ferritin (50×) | 1.4 | 4 | Backward | 115.4 | 1,102 | Permanently Clogged |
| Ferritin (4×) | 17 | 4 | Backward | 135 | 63,119 | Open |
| Total Events | | | | | 2,877,759 | |

**Table S1:** A representative set of pores and proteins used in the study. Color-coded for ease of identification.

### Section 6: Zeta Potential

The surface charge density of a surface ($\sigma_p$) can be calculated using Grahame's equation, $\sigma_p = \frac{2\epsilon_r\epsilon_0\kappa}{\beta e} sinh\left(\frac{\beta e\phi_p}{2}\right)$ where $\epsilon_r\epsilon_0$, $\kappa^{-1}$, $\beta$, $e$ and $\phi_p$ are the permittivity of the solution, Debye screening length, the inverse of thermal energy, elementary charge, and the diffuse layer potential respectively.[1-3] For a negatively charged surface with a dissociation constant of $pK$, one could express $\phi_p$ as, $\phi_p = \frac{1}{\beta e} ln \frac{-\sigma_p}{e\Gamma+\sigma_p} - (pH - pK)\frac{ln10}{\beta e} - \frac{\sigma_p}{C}$ where $\Gamma$, $C$ are the total surface density of surface chargeable groups, and Stern layer capacitance. For all calculations, 0.3 F/m$^2$, 8.5, 77.75, $8.854 \times 10^{-12}$ F/m$^2$ and $8\times10^{18}$ m$^{-2}$ were used for $C$, $pK_a$, $\epsilon$, $\epsilon_0$ and $\Gamma$ respectively.[4] For lower potentials, Grahame's equation can be approximated to (assuming $\phi_p \approx \zeta_p$ where $\zeta_p$ is the zeta potential of the nanopore surface)[1, 5] $\zeta_p = \frac{\sigma_p}{\epsilon_r\epsilon_0}\kappa^{-1}$.

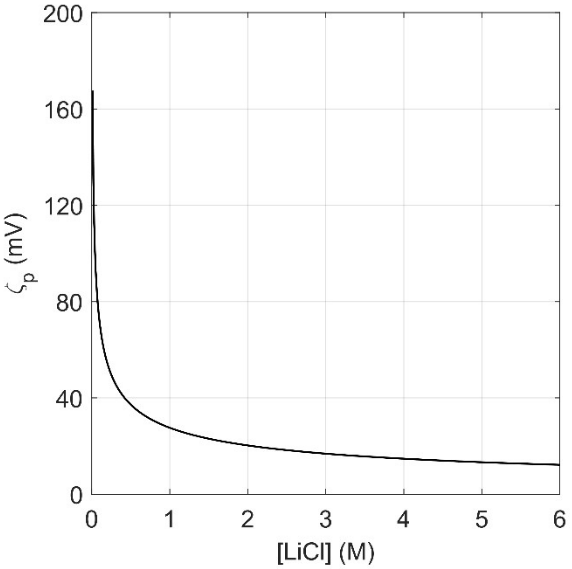

**Figure S6:** Absolute value of the zeta potential of the pore surface ($\zeta_p$) calculated as shown in this section versus the concentration of LiCl.

### Section 7: COMSOL Model

The model was built using COMSOL Multiphysics 5.6 using a 2D axisymmetric assumption. The geometry is shown in Figure S7 wherein an angled rectangle with rounded edges is used as the nanopipette wall and separates the two fluidic half cells of the nanopore system. COMSOL simultaneously solves the Nernst-Planck, Poisson's, and Navier-Stokes equations under stationary (steady state) conditions. The modules used within COMSOL which utilize and simultaneously solve these equations include: Transport of Dilute Species, Electrostatics, and Laminar Flow. The boundary conditions for the pore wall designating a surface charge (σ) of -0.02 C/m$^2$ and an electroosmotic flow boundary condition based on the zeta potential (ζ). The outer circle of the fluidic cell representing the fluidic "bath" was set as an outlet with a pressure designation of 1 atmosphere and electrostatic ground. The boundary inside the nanopipette was designated as having an electric potential equal to the applied voltage and a fluidic inlet with the same pressure as the outlet (1 atmosphere). Under no voltage bias, there was not pressure driven flow generated by these boundary conditions. Incompressible liquid and charge conservation were used for the internal domains of the model. The default parameters used throughout the modeling process (unless otherwise noted) were $d_p$ = 10 nm, μ (electrophoretic mobility) = $1\times10^{-8}$ m$^2$/Vs, V = -600 mV and ζ=-20 mV.

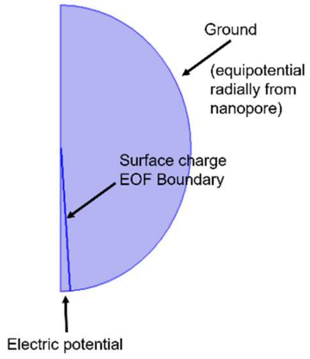

**Figure S7:** Schematic of the 2D axisymmetric model used to numerically solve for the fluid flow and electric potential within the nanopipette. The outer sphere diameter was 1 μm and the half cone angle of the pipette was set to 7 degrees.

### References

1. Arjmandi, N.; Van Roy, W.; Lagae, L.; Borghs, G., Measuring the electric charge and zeta potential of nanometer-sized objects using pyramidal-shaped nanopores. *Analytical Chemistry* **2012,** *84* (20), 8490-8496.
2. Smeets, R. M.; Keyser, U. F.; Krapf, D.; Wu, M.-Y.; Dekker, N. H.; Dekker, C., Salt dependence of ion transport and DNA translocation through solid-state nanopores. *Nano Letters* **2006,** *6* (1), 89-95.
3. Frament, C. M.; Bandara, N.; Dwyer, J. R., Nanopore surface coating delivers nanopore size and shape through conductance-based sizing. *ACS Applied Materials & Interfaces* **2013,** *5* (19), 9330-9337.
4. van der Heyden, F. H.; Stein, D.; Dekker, C., Streaming currents in a single nanofluidic channel. *Physical Review Letters* **2005,** *95* (11), 116104.
5. Kejian, D.; Weimin, S.; Haiyan, Z.; Xianglei, P.; Honggang, H., Dependence of zeta potential on polyelectrolyte moving through a solid-state nanopore. *Applied Physics Letters* **2009,** *94* (1), 014101.
